# Backbone and Sidechain ^1^H, ^15^N and ^13^C Resonance Assignments of a Multidrug Efflux Membrane Protein using Solution and Solid-State NMR

**DOI:** 10.1101/2025.01.07.631770

**Authors:** Benjamin D. Harding, Ashley Hiett, Marco Tonelli, Songlin Wang, Chad M. Rienstra, Katherine A. Henzler-Wildman

**Affiliations:** Biophysics Graduate Program, University of Wisconsin-Madison, Madison, WI, 53706 USA; Department of Biochemistry, University of Wisconsin-Madison, Madison, WI, 53706 USA; National Magnetic Resonance Facility at Madison, University of Wisconsin-Madison, Madison, WI, 53706 USA; Morgridge Institute for Discovery, University of Wisconsin-Madison, Madison, WI, 53706 USA

**Keywords:** EmrE, membrane protein, antibiotic resistance, solid-state NMR, NMR spectroscopy, magic-angle spinning

## Abstract

EmrE is a bacterial membrane-embedded multidrug transporter that functions as an asymmetric homodimer. EmrE is implicated in antibiotic resistance, but is now known to confer either resistance or susceptibility depending on the identity of the small molecule substrate. Here, we report both solution- and solid-state NMR assignments of S64V-EmrE at pH 5.8, below the pKa of critical residues E14 and H110. This includes ^1^H, ^15^N, and ^13^C resonance assignments of the backbone, methyl groups (isoleucine, leucine, valine, threonine and alanine) from solution NMR experiments in bicelles, and backbone and side-chain assignments from solid-state NMR ^13^C-detected experiments in liposomes.

## Biological Context

Small multidrug resistance (SMR) transporters are a family of membrane embedded efflux pumps found throughout the bacterial kingdom that contribute to antibiotic resistance through efflux of toxic compounds (Paulsen et al. 1996). EmrE is an SMR transporter found in the inner membrane of *E. coli* and is a model system to study the minimal requirements of proton-coupled transport (Yerushalmi 2000). The foundational structural unit of EmrE is an antiparallel homodimer (Morrison et al. 2012; Tate et al. 2001). Each subunit consists of 110 amino acids with four transmembrane (TM) helices connected by interhelical loops and capped by a C-terminal tail. The binding pocket is made up of TM helices 1-3 from each subunit and consists of essential binding residues E14_A_ and E14_B_ along with bulky aromatic residues. The TM 4 helices from each subunit interact to form EmrE’s dimerization domain. Biochemical and biophysical data also suggest the interhelical loops and C-terminal residues play critical roles in regulating EmrE’s promiscuous multidrug recognition and transport pathways (Glaubitz et al. 2000; Spreacker et al. 2022; Thomas et al. 2018).

Here, we use ^13^C-detected solid-state NMR (SSNMR) to assign backbone and sidechain chemical shifts and solution NMR to assign backbone and methyl groups of EmrE mutant S64V at pH 5.8 (BMRB ID 51024). The S64V point mutant is located within the third TM helix and lowers the rate of alternating access while retaining binding affinity to substrate (Wu et al. 2019). Chemical shifts between solution and solid state agree well, indicating the structure of EmrE is preserved over a large temperature range and across different membrane mimetics, since solution data was recorded at 45 °C in q=0.33 DMPC/DHPC isotropic bicelles and solid-state data was recorded both above and below the phase transition for protein in POPC liposomes.

## Methods and Experiments

### Protein Expression and purification

S64V-EmrE was expressed from a pET15b plasmid in *E. coli* BL21(DE3) as previously described (Morrison et al. 2012; Wu et al. 2019). Protein for solid-state NMR samples was expressed in *E. coli* grown in ^1^H/^15^N-M9 minimal media supplemented with 2.5 g/L D-glucose-^13^C_6_ or 2.5 g/L 1,3-^13^C_2_-glycerol as the sole carbon source. Protein for solution NMR backbone walk experiments was expressed in *E. coli* grown in ^2^H/^15^N-M9 minimal media supplemented with 2.5 g/L D-Glucose-^13^C_6_,1,2,3,4,5,6,6-d_7_. Protein for solution NMR methyl experiments was expressed in *E. coli* grown in ^2^H/^15^N-M9 minimal media supplemented with 2.5 g/L D-Glucose-1,2,3,4,5,6,6-d_7_ and addition of various methyl precursors one hour before induction according to previously established protocols (Kurauskas et al. 2017): (1) ^2^H/^15^N ILVTA-^13^CH_3_-methyl sample for NOESY experiments at 900 MHz with labelled Ile Cδ1, Leu CA, CB, CG, and Cδ1 or Cδ2, Val CA, CB, and Cγ1 or Cγ2, Thr Cγ2, Ala CB, prepared using 60 mg/L 2-ketobutyric Acid-4-13C,3,3-dz Sodium Salt Hydrate, 120 mg/L Alpha-Ketoisovaleric Acid Sodium Salt (1,2,3,4-13C4, 99%; 3,4,4,4-D4, 97-98%), 50 mg/L Threonine (4-13C,97%, 2,3-D2, 97-98%), 600 mg/L L-Alanine (3-13C, 99%, 2-D 96%), and 2500 mg/L Succinic Acid, D4 98%. (2) Fully ^13^C LV sample for TOCSY experiments at 750 MHz and short mixing-time NOESY experiments at 600 MHz with labelled Leu CA, CB, CG, Cδ1 and Cδ2, Val CA, CB, Cγ1 and Cγ2, prepared using −120 mg/L alpha-Ketoisovaleric acid, sodium salt (¹³C₅, 98%; 3-D₁, 98%). (3) ^2^H/^15^N ILVT-^13^CH_3_-methyl sample for NOESY experiments at 1.1 GHz with labelled Ile Cδ1, Leu Cδ1 or Cδ2, Val Cγ1 or Cγ2, Thr Cγ2, prepared using 200 mg/L Alpha-Ketoisovaleric acid, sodium salt (3-methyl-¹³C, 99%; 3,4,4,4-D₄, 98%), 60 mg/L alpha-Ketobutyric acid, sodium salt (methyl-¹³C, 99%; 3,3-D₂, 98%), and 50 mg/L L-Threonine (4-¹³C, 97%; 2,3-D₂, 96-98%). In samples with labeled Thr Cγ2, threonine contained ^14^N in its amide group, causing threonine residues to become invisible in amide NMR experiments.

Purification for all samples was conducted as previously described (Morrison and Henzler-Wildman 2012). Briefly, protein was purified in *n*-decyl-β-maltoside (DM) with metal affinity chromatography followed by size exclusion chromatography. Fractions containing purified EmrE in DM were added to POPC (solid-state samples) or DMPC (solution samples) lipid suspension at the protein to lipid ratio listed below and incubated at 25 °C for 20 min. Detergent was removed with Amberlite (Supelco) hydrophobic beads overnight and liposomes collected by ultracentrifugation. Solid state NMR samples were prepared by reconstituting purified S64V-EmrE in DM into POPC proteoliposomes at a 25:1 lipid/EmrE monomer ratio as described in (Harding et al. 2024). Excess water was removed from the concentrated proteoliposomes by freezing the sample tube in liquid nitrogen, immediately microcentrifuging at 15,000*g* for 30 s, then decanting. Solution NMR samples were prepared by reconstituting S64V-EmrE into DMPC (14:0 PC-d54) liposomes at a 75:1 lipid: EmrE monomer molar ratio in a similar process to (Morrison et al. 2012). Isotropic bicelles (q=0.33) were then formed by adding deuterated DHPC (06:0 PC-d22) to the concentrated proteoliposomes and freeze/thawing three times.

## Solution NMR Spectroscopy

All solution NMR samples had a final concentration of 0.5-1 mM EmrE reconstituted into isotropic bicelles in 20 mM NaCl, 20 mM MES buffer, pH 5.8 with 10% D_2_O, 2 mM tris(2-carboxyethyl)phosphine hydrochloride (TCEP), 0.05% NaN_3_, and 50 µM 2,2-dimethyl-2-silapentanesulfonic acid (DSS) as an internal standard. Solution NMR samples were transferred to 3 mm or 5 mm Shegemi tubes with plungers for NMR experiments. Experiments were all conducted at the National Magnetic Resonance Facility at Madison (NMRFAM) at 318 K. All 3D and 4D experiments were collected with 25% non-uniform sampling (NUS) (Kazimierczuk and Orekhov 2011; Mayzel et al. 2014). Spectra were processed with NMRPipe (Delaglio et al. 1995) using IST reconstruction (Stern et al. 2007) or SMILE reconstruction (Ying et al. 2017) of NUS experiments and analyzed using CCPNMR v3.0 (Skinner et al. 2016). All experiments used in assigning solution NMR resonances of pH 5.8 S64V-EmrE are summarized with relevant experimental details in Table I.

**Table I:**
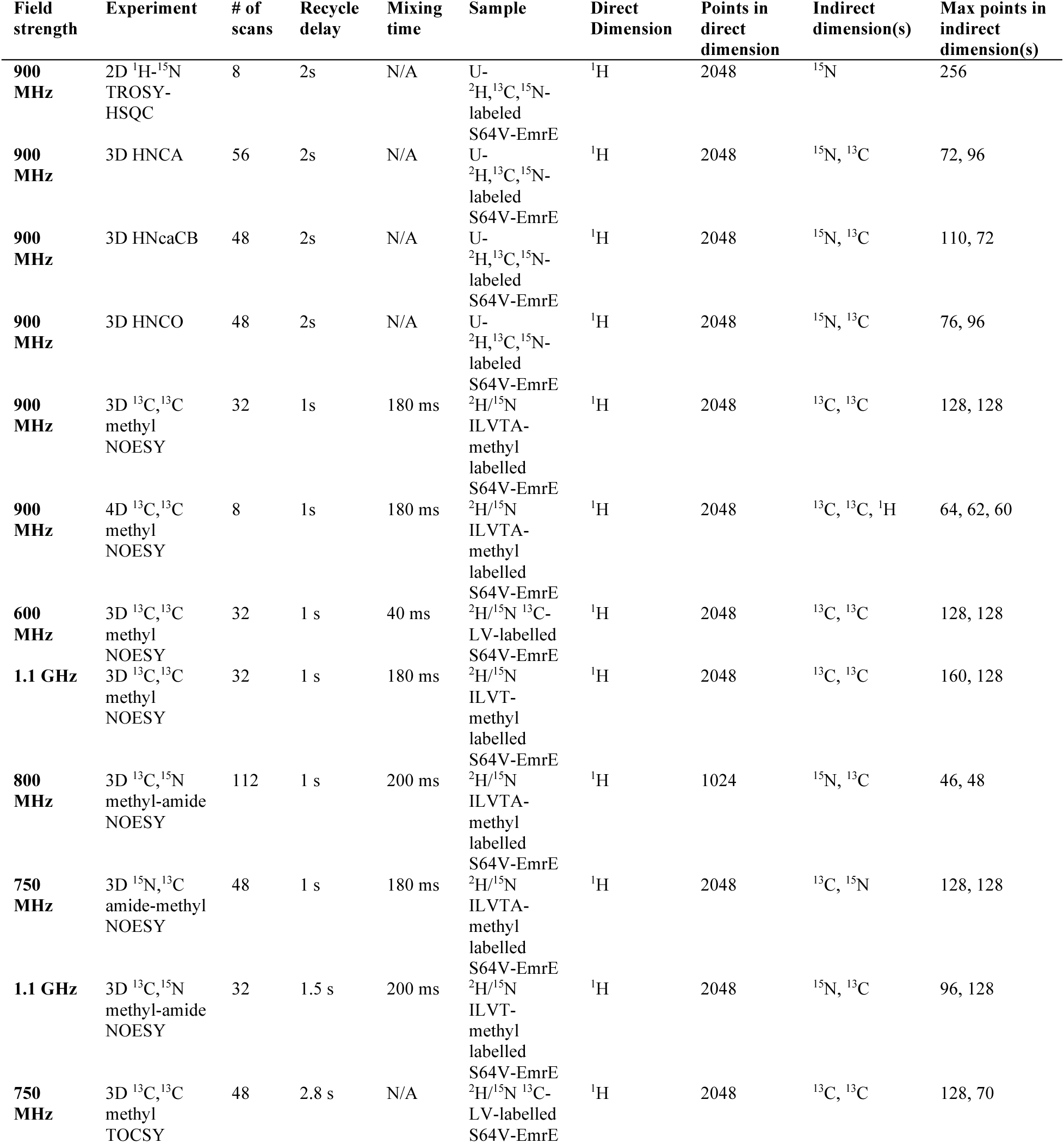
Summary of solution NMR experiments and experimental parameters used in assigning resonances of pH 5.8 S64V EmrE in bicelles.

| Field strength | Experiment | # of scans | Recycle delay | Mixing time | Sample | Direct Dimension | Points in direct dimension | Indirect dimension(s) | Max points in indirect dimension(s) |
| --- | --- | --- | --- | --- | --- | --- | --- | --- | --- |
| 900 MHz | 2D $^1\text{H}$-$^{15}\text{N}$ TROSY-HSQC | 8 | 2s | N/A | U-$^2\text{H}$, $^{13}\text{C}$, $^{15}\text{N}$-labeled S64V-EmrE | $^1\text{H}$ | 2048 | $^{15}\text{N}$ | 256 |
| 900 MHz | 3D HNCA | 56 | 2s | N/A | U-$^2\text{H}$, $^{13}\text{C}$, $^{15}\text{N}$-labeled S64V-EmrE | $^1\text{H}$ | 2048 | $^{15}\text{N}$, $^{13}\text{C}$ | 72, 96 |
| 900 MHz | 3D HNcaCB | 48 | 2s | N/A | U-$^2\text{H}$, $^{13}\text{C}$, $^{15}\text{N}$-labeled S64V-EmrE | $^1\text{H}$ | 2048 | $^{15}\text{N}$, $^{13}\text{C}$ | 110, 72 |
| 900 MHz | 3D HNCO | 48 | 2s | N/A | U-$^2\text{H}$, $^{13}\text{C}$, $^{15}\text{N}$-labeled S64V-EmrE | $^1\text{H}$ | 2048 | $^{15}\text{N}$, $^{13}\text{C}$ | 76, 96 |
| 900 MHz | 3D $^{13}\text{C}$, $^{13}\text{C}$ methyl NOESY | 32 | 1s | 180 ms | $^2\text{H}/^{15}\text{N}$ ILVTA-methyl labelled S64V-EmrE | $^1\text{H}$ | 2048 | $^{13}\text{C}$, $^{13}\text{C}$ | 128, 128 |
| 900 MHz | 4D $^{13}\text{C}$, $^{13}\text{C}$ methyl NOESY | 8 | 1s | 180 ms | $^2\text{H}/^{15}\text{N}$ ILVTA-methyl labelled S64V-EmrE | $^1\text{H}$ | 2048 | $^{13}\text{C}$, $^{13}\text{C}$, $^1\text{H}$ | 64, 62, 60 |
| 600 MHz | 3D $^{13}\text{C}$, $^{13}\text{C}$ methyl NOESY | 32 | 1 s | 40 ms | $^2\text{H}/^{15}\text{N}$ $^{13}\text{C}$-LV-labelled S64V-EmrE | $^1\text{H}$ | 2048 | $^{13}\text{C}$, $^{13}\text{C}$ | 128, 128 |
| 1.1 GHz | 3D $^{13}\text{C}$, $^{13}\text{C}$ methyl NOESY | 32 | 1 s | 180 ms | $^2\text{H}/^{15}\text{N}$ ILVT-methyl labelled S64V-EmrE | $^1\text{H}$ | 2048 | $^{13}\text{C}$, $^{13}\text{C}$ | 160, 128 |
| 800 MHz | 3D $^{13}\text{C}$, $^{15}\text{N}$ methyl-amide NOESY | 112 | 1 s | 200 ms | $^2\text{H}/^{15}\text{N}$ ILVTA-methyl labelled S64V-EmrE | $^1\text{H}$ | 1024 | $^{15}\text{N}$, $^{13}\text{C}$ | 46, 48 |
| 750 MHz | 3D $^{15}\text{N}$, $^{13}\text{C}$ amide-methyl NOESY | 48 | 1 s | 180 ms | $^2\text{H}/^{15}\text{N}$ ILVTA-methyl labelled S64V-EmrE | $^1\text{H}$ | 2048 | $^{13}\text{C}$, $^{15}\text{N}$ | 128, 128 |
| 1.1 GHz | 3D $^{13}\text{C}$, $^{15}\text{N}$ methyl-amide NOESY | 32 | 1.5 s | 200 ms | $^2\text{H}/^{15}\text{N}$ ILVT-methyl labelled S64V-EmrE | $^1\text{H}$ | 2048 | $^{15}\text{N}$, $^{13}\text{C}$ | 96, 128 |
| 750 MHz | 3D $^{13}\text{C}$, $^{13}\text{C}$ methyl TOCSY | 48 | 2.8 s | N/A | $^2\text{H}/^{15}\text{N}$ $^{13}\text{C}$-LV-labelled S64V-EmrE | $^1\text{H}$ | 2048 | $^{13}\text{C}$, $^{13}\text{C}$ | 128, 70 |

### Backbone Walk Experiments

Backbone double and triple resonance experiments were acquired on a U-^2^H,^13^C,^15^N-labeled S64V-EmrE sample in q=0.33 DMPC/DHPC isotropic bicelles at pH 5.8. 2D ^1^H-^15^N TROSY-HSQC, and 3D HNCA, HNcaCB, and HNCO (Michael Sattler 1999) were acquired on a Bruker Avance III HD spectrometer at 900 MHz equipped with a 5 mm TCI cryoprobe.

Backbone assignments of S64V-EmrE at pH 5.0 in in DMPC/DHPC isotropic bicelles were previously performed using HNCO, HNcaCO and HNCA, HNcoCA, paired experiments (BMRB ID 51024) (Spreacker et al. 2022). The close correspondence of the experimental conditions and backbone chemical shifts allowed complete assignment confirmation with a reduced set of 3D spectra under the current conditions at pH 5.8. Notably, the loop residues R82-D84 and tail residues R106-H110 of both chains appear to have two distinct sets of ^15^N and ^1^H resonances, one with high peak intensity and one with low peak intensity, whose CA and CB chemical shifts match each other identically.

### Methyl-Methyl NOESY Experiments

3D and 4D ^13^C,^13^C methyl NOESY experiments (Catherine Zwahlen 1998) were performed on the ^2^H/^15^N ILVTA-methyl labelled S64V-EmrE sample using a Bruker Avance III HD spectrometer at 900 MHz equipped with a 5 mm TCI cryoprobe. A 3D ^13^C,^13^C methyl NOESY experiment was acquired for the ^2^H/^15^N geminal LV-labelled S64V-EmrE sample using a Bruker Avance III HD spectrometer at 600 MHz equipped with a 5 mm TCI cryoprobe. A 3D ^13^C,^13^C methyl NOESY experiment was acquired for the ^2^H/^15^N ILVT-methyl labelled S64V-EmrE sample using a Bruker NEO spectrometer at 1.1 GHz equipped with a 3 mm TCI cryoprobe.

### Methyl-Amide NOESY Experiments

3D ^13^C,^15^N methyl-amide NOESY experiment was carried out on the ^2^H/^15^N ILVTA-methyl labelled S64V-EmrE sample using a Varian VNMRS DD spectrometer at 800 MHz equipped with a 5 mm cold probe. A 3D ^15^N,^13^C amide-methyl NOESY experiment was conducted on the same sample using a Bruker Avance III HD spectrometer at 750 MHz equipped with a 5 mm TXI cryoprobe. A 3D ^13^C,^15^N methyl-amide NOESY experiment was acquired on the ^2^H/^15^N ILVT-methyl labelled S64V-EmrE sample using a Bruker NEO spectrometer at 1.1 GHz equipped with a 3 mm TCI cryoprobe.

### Methyl TOCSY Experiments

The 3D ^13^C,^13^C methyl TOCSY experiment (Uhrin et al. 2000) was acquired with the ^2^H/^15^N ^13^C-LV-labelled S64V-EmrE sample using a Bruker Avance III HD spectrometer at 750 MHz equipped with a 5 mm TXI cryoprobe.

## Solid-State NMR Spectroscopy

Samples of uniformly ^13^C-^15^N labeled and sparsely labeled EmrE grown with 1,3-glycerol (Cambridge Isotope Laboratories, Andover MA) were reconstituted into POPC liposomes as described above (Castellani et al. 2002). These samples were packed into PhoenixNMR (Loveland, CO) 3.2 mm magic-angle spinning (MAS) rotor using custom-designed rotor packing devices as previously described (Olson et al. 2024). Uniformly-labeled EmrE in proteoliposomes were also packed into a Phoenix 1.6 mm MAS rotor in a similar manner. The total amount of protein packed into the 3.2 mm rotors was 7.5 mg, and 2.5 mg in the 1.6 mm rotor, based on a 60% hydrated sample and a lipid-to-protein molar ratio of 25:1. This formulation exhibits a phase transition at −15 ± 5 °C. Spectra obtained both above and below the phase transition exhibit high resolution and sensitivity, although the sensitivity and experimental stability are slightly better for experiments conducted below the phase transition when requiring multiple cross polarization (CP) steps. Spectra were referenced to DSS, using adamantane as a secondary external standard with the downfield ^13^C signal at 40.48 ppm (Morcombe and Zilm 2003).

### 1.1 GHz Experiments

SSNMR experiments at 1.1 GHz were performed at NMRFAM using a Bruker NEO spectrometer equipped with a Blackfox 1.6 mm triple resonance probe in HCN mode. The variable temperature gas set point was −10 °C with 20 lpm flow at an MAS rate of 25 kHz, resulting in a sample temperature of 10 ± 5 °C. Indirect dimensions were sampled with States-TPPI phase cycling (Dominique Marion 1989). Pulse widths for ^1^H, ^13^C, and ^15^N were 2.8, 2.5, and 7.4 μs, respectively. All experiments utilized 100 kHz SPINAL-64 ^1^H decoupling during evolution and acquisition times.

#### ^13^C-^13^C and ^15^N-^13^C 2D Experiments

The 2D ^13^C-^13^C spectrum was collected with 1.5 ms CP contact time with nutation frequencies ωC = 108 kHz and a downward tangent ramp (Hediger et al. 1994; S. Hediger 1995) on ^1^H with an average RF amplitude of 82 kHz. Homonuclear recoupling used CORDxy4, 50 ms. (Hou et al. 2013). The maximum evolution time (t_1_) was 8.0 ms (5 µs x 1600 total points), and the t_2_ acquisition time was 20.5 ms (5 µs x 4096 total points). The recycle delay was 1.5 s with 16 scans per row. The total experimental time was 11.3 hours.

The 2D NCA experiment was performed with the ^13^C carrier frequency at 55.08 ppm and 117.34 ppm for ^15^N. The ^1^H to ^15^N CP contact time was 5 ms with nutation frequencies ωN = 40 kHz and a downward tangent ramp on ^1^H with an average RF amplitude of 59 kHz. The NCA SPECIFIC CP (Baldus et al. 1998) used a double quantum (DQ) n=1 condition with ωN = 10 kHz and an upward tangent ramp on ^13^C with an average RF amplitude of 15 kHz. The contact time was 5 ms with 100 kHz continuous wave (CW) decoupling. The t_1_ maximum evolution time was 11.5 ms (80 μs x 144 total points) and the t_2_ acquisition time was 20.5 ms (5 μs x 4096 total points). LOW-BASHD decoupling was implemented in the direct dimension using τ_dec_=3.2 ms and 72.5 µs Gaussian π-pulses with a maximum amplitude of ωC = 17 kHz and carrier frequency at 175 ppm. (Struppe et al. 2013) The recycle delay was 1.5 s with 256 scans per row. The total experimental time was 15.8 hr.

The 2D NCO experiment was performed with the ^13^C carrier frequency at 174.92 ppm and 117.35 ppm for ^15^N. The ^1^H to ^15^N CP contact time was 1 ms with nutation frequencies ωN = 38 kHz and a downward tangent ramp on ^1^H with an average RF amplitude of 60 kHz. The NCO SPECIFIC CP used a double quantum (DQ) n=1 condition with nutation frequencies ωN = 16 kHz and an upward tangent ramp on ^13^C with an average RF amplitude of 10 kHz. The contact time was 3 ms with CW decoupling on ^1^H (100 kHz). The t_1_ maximum evolution time was 11.5 ms (80 μs x 144 total points) and the t_2_ acquisition time was 20.5 ms (5 μs x 4096 total points). LOW-BASHD decoupling was implemented in the direct dimension using τ_dec_=3.2 ms and 72.5 µs Gaussian π-pulses with a maximum amplitude of ωC = 17 kHz and carrier frequency at 175 ppm. (Struppe et al. 2013) The recycle delay was 1.5 s with 256 scans per row. The total experimental time was 15.8 hr.

#### ^15^N-^13^C-^13^C and ^13^C-^15^N-^13^C 3D Experiments

In all 3D experiments, LOW-BASHD decoupling was implemented in the direct dimension using τ_dec_=3.2 ms and 72.5 µs Gaussian π-pulses with a maximum amplitude of ωC = 17 kHz and carrier frequency at 175 ppm. (Struppe et al. 2013) All 3D experiments were collected using non-uniform sampling (NUS) using a 25% randomly sampled Poisson gap schedule generated from an in-house script. (Hyberts et al. 2012)

The 3D NCACO experiment was performed with the ^13^C carrier frequency set to 54.92 ppm and 117.35 for ^15^N. The ^1^H to ^15^N CP contact time was 5 ms with nutation frequencies ωN = 40 kHz and a downward tangent ramp on ^1^H with an average RF amplitude of 59 kHz. The NCA SPECIFIC CP used a double quantum (DQ) n=1 condition with ωN = 10 kHz and an upward tangent ramp on ^13^C with an average RF amplitude of 15 kHz. The contact time was 5 ms with 100 kHz continuous wave (CW) decoupling. The t_1_ maximum evolution time was 10.8 ms (100 μs x 108 total points), the t_2_ maximum evolution time was 4.8 ms (40 μs x 120 total points), and the t_3_ acquisition time was 20.5 ms (5 μs x 4096 total points). The recycle delay was 1.5 s and each spectrum was collected with 16 scans per row. Five replicate spectra were collected and co-added in the time domain. The total experimental time was 112 hr.

The 3D NCOCA experiment was performed with the ^13^C carrier frequency at 175.08 ppm and 117.35 ppm for ^15^N. The ^1^H to ^15^N CP contact time was 1 ms with nutation frequencies ωN = 38 kHz and a downward tangent ramp on ^1^H with an average RF amplitude of 60 kHz. The NCO SPECIFIC CP used a double quantum (DQ) n=1 condition with nutation frequencies ωN = 16 kHz and an upward tangent ramp on ^13^C with an average RF amplitude of 10 kHz. The contact time was 3 ms with CW decoupling on ^1^H (100 kHz). The ^13^C-^13^C polarization transfer utilized CORDxy4 (50 ms) with a ^1^H RF amplitude of 25 kHz. The t_1_ maximum evolution time was 10.8 ms (100 μs x 108 total points), the t_2_ maximum evolution time was 5.1 ms (80 μs x 64 total points), and the t_3_ acquisition time was 20.5 ms (5 μs x 4096 total points). The recycle delay was 1.5 s and each spectrum was collected with 16 scans per row. Four replicate spectra were collected and co-added in the time domain. The total experimental time was 95 hr.

The 3D CANCO experiment was performed with the ^13^CA, ^13^C’,and ^15^N carrier frequencies set to 55, 174.92, and 117.35 ppm, respectively. CP from ^1^H to ^13^CA used nutation frequencies of ωC = 108 kHz and an upward tangent ramp on the ^1^H channel with an average RF amplitude of 82 kHz at a contact time of 0.5 ms. CP from ^13^CA to ^15^N used nutation frequencies of ωN = 10 kHz with an upward tangent ramp on the ^13^C channel with an average RF amplitude of 16 kHz at a contact time of 3 ms with 100 kHz CW decoupling. CP from ^15^N to ^13^C’ used nutation frequencies of ωN = 16 kHz and an upward tangent ramp on the ^13^C channel with an average RF amplitude of 10 kHz at a contact time of 3 ms with 100 kHz ^1^H CW decoupling. The maximum t_1_ evolution time was 4.8 ms (40 μs x 120 total points), the t_2_ maximum evolution time was 10.8 ms (100 μs x 108 total points), and the t_3_ acquisition time was 20.5 ms (5 μs x 4096 total points). The recycle delay was 1.5 s and each spectrum was collected with 16 scans per row. Five replicate spectra were collected and co-added in the time domain. The total experimental time was 112 hr.

### 750 MHz Experiments

Spectra collected at 750 MHz utilized a Varian 17.6 T wide bore magnet, VNMRS console, custom-configured amplifiers as described in (Harding et al. 2024) and a Varian Balun 3.2 mm probe in double resonance HC mode for maximum ^13^C sensitivity. (Han et al. 2024) Data sets were collected over a range of temperatures to determine that the optimal sensitivity was observed below the lipid phase transition at a sample temperature −15 ± 5 °C and MAS rate 12.5 kHz. Sparsely labeled EmrE grown with 1,3-glycerol was collected above the phase transition at sample temperature of −13 ± 5 °C where resolution was optimal. The ^1^H and ^13^C pulse widths were 2.6 and 2 μs in the double resonance configuration for CC and CCC experiments in spectra collected above and below the phase transition. For HCN mode experiments, ^1^H, ^13^C, and ^15^N pulse widths were 2.7, 3, and 4.8 μs, respectively. In all cases, and SPINAL-64 decoupling was optimized at 90 kHz with a 5.4 μs π pulse (Comellas et al. 2011).

#### ^13^C-^13^C 2D and ^13^C-^13^C-^13^C 3D Experiments

##### Uniformly ^13^C-^15^N labeled EmrE spectra collected below the phase transition

CP from ^1^H to ^13^C used nutation frequencies of ωH = 80 kHz and ωC = 97 kHz with a tangent ramp on the ^1^H channel with Δ and β parameters of 9.6 kHz and a contact time of 1 ms, where the Δ parameter defines the magnitude of the difference in the RF field and the β parameter defines the curvature of the ramp. For ^13^C-^13^C 2D experiments with 50, 125, and 250 ms dipolar assisted rotational resonance (DARR) mixing times were signal averaged for a total of 2, 10 and 10 hours, in blocks of 4, 8, and 8 scans per row repeated 4, 2 and 2 times. The t_1_ maximum evolution time was 10.2 ms (20 μs x 512 complex points) and the t_2_ acquisition time was 15 ms (10 μs x 1500 complex points).

Two 3D ^13^C-^13^C-^13^C experiments were carried out with a two-step phase cycle and collected with DARR mixing times of t_mix,1_ = t_mix,2_ = 50 ms and another with t_mix,1_ = 50 ms and t_mix,2_ = 250 ms. The t_1_ and t_2_ maximum evolution times were both 7.7 ms (20 μs x 384 complex points) and the t_3_ acquisition time was 15 ms (10 μs x 1500 complex points). Both spectra were collected using NUS where the schedule was constructed from the website of Wagner and co-workers (http://gwagner.med.harvard.edu/intranet/hmsIST/gensched_new.html) as a 384 x 384 point randomly sampled sine squared weighted grid with 25% coverage.

##### Sparsely labeled EmrE above the phase transition

CP from ^1^H to ^13^C used nutation frequencies of ωH = 73 kHz and ωC = 58 kHz with a tangent ramp on the ^13^C channel (Δ = 14 kHz and β = 2 kHz) with a contact time of 700 μs. 2D ^13^C-^13^C experiments with 150 and 600 DARR mixing times were signal averaged for a total of 40 and 240 hr, in blocks of 5 and 24 and 16 scans per row. The t_1_ maximum evolution time was 10.2 ms (20 μs x 512 complex points) and the t_2_ acquisition time was 15 ms (10 μs x 1500 complex points).

A 3D ^13^C-^13^C-^13^C experiment was carried out with a two-step phase cycle and collected with DARR mixing times of t_mix,1_ = 150 ms and t_mix,2_ = 600 ms. The t_1_ and t_2_ maximum evolution times were both 7.7 ms (20 μs x 384 complex points) and the t_3_ acquisition time was 15 ms (10 μs x 1500 complex points). The spectrum was collected using NUS where the schedule was constructed from the website of Wagner and co-workers as a 384 x 384 point randomly sampled sine squared weighted grid with 25% coverage.

#### ^15^N-^13^C-^13^C and ^13^C-^15^N-^13^C 3D Experiments

Triple resonance ^15^N-^13^C-^13^C and ^13^C-^15^N-^13^C experiments were collected at 17.6 T equipped with a 3.2 mm Balun probe in HCN mode with the same speed limited Varian rotor packed with uniformly ^13^C-^15^N labeled EmrE spinning at 12.5 kHz at sample temperature below the phase transition (−20 ± 5 °C).

The 3D NCACX experiment was collected with the ^13^C and ^15^N carrier frequencies at 59.94 and 98.03 ppm, respectively. CP from ^1^H to ^15^N used nutation frequencies of ωH = 71 kHz and ωN = 51 kHz with a tangent ramp on ^1^H with Δ and β parameters of 9.3 kHz with a contact time of 1.2 ms. CP from ^15^N to ^13^C used nutation frequencies of ^15^N to ^13^C was achieved with ωN = 34.5 kHz and ωC = 18.5 kHz with a tangent ramp on the ^13^C channel with Δ = −1 kHz and β = 1 kHz with a contact time of 6 ms. A band-selective rSNOB soft pulse (520 μs) was employed at a carrier frequency of 60 ppm, and DARR (t_mix_ = 50 ms) was used for homonuclear recoupling. The t_1_ maximum evolution time was 20.5 ms (80 μs x 256 complex points), the t_2_ maximum evolution time was 6.4 ms (80 μs x 80 complex points), and the t_3_ acquisition time was 15 ms (10 μs x 1500 complex points). The spectrum was collected using NUS and a 50% randomly sampled Poisson-gap schedule generated on the website of Gerhard and co-workers with maximum evolution times 80 x 256 hypercomplex points in the ^13^C and ^15^N dimensions, respectively. The spectrum was collected with a 1.5 s recycle delay and four scans per row. The total experimental time was 73 hr.

The 3D NCOCX was collected with the ^13^C and ^15^N carrier frequencies at 175.14 and 110.13 ppm, respectively. CP from ^1^H to ^15^N used nutation frequencies of ωH = 83 kHz and ωN = 51 kHz with a tangential ramp on the ^1^H channel with Δ = 14 kHz and β = 5 kHz with a contact time of 1.6 ms. CP from ^15^N to ^13^C used nutation frequencies of ωN = 32 kHz and ωC = 44 kHz with a tangential ramp on the ^13^C channel with Δ = −2 kHz and β = 2 kHz with a contact time of 6 ms. DARR mixing (t_mix_ = 50 ms) was employed. The t_1_ maximum evolution time was 16 ms (160 μs x 100 complex points), the t_2_ maximum evolution time was 12.8 ms (160 μs x 80 complex points), and the t_3_ acquisition time was 15 ms (10 μs x 1500 complex points). The spectrum was collected using NUS and a 50% randomly sampled Poisson-gap schedule generated on the website of Gerhard and co-workers with maximum evolution times 80 x 100 hypercomplex points in the ^13^C and ^15^N dimensions, respectively. The NCOCX experiment was repeated three times, and each spectrum took 32 hr to collect. Each spectrum was collected with a recycle delay of 1.75 s and four scans per row. The total experimental time was 96 hr.

The 3D CANCO experiment was collected under the conditions described in (Harding et al. 2024).

## Extent of Assignments and Data Deposition

### Protein Backbone Assignments

The 2D NCA and HSQC spectra are unique fingerprint spectra in SSNMR and solution NMR respectively and serve as a roadmap when performing backbone assignments on EmrE. Addition of the ^15^N dimension in SSNMR breaks chemical shift degeneracy of repetitive hydrophobic residues with highly degenerate ^13^C backbone chemical shifts, notably valine, isoleucine, leucine, and alanine, which comprise 41.8% of the amino acid sequence in EmrE. The 2D NCA (Fig. 1A) shows exquisite resolution and sensitivity, allowing for unambiguous identification of identical, but chemically inequivalent, amino acids in chain A and chain B of the asymmetric EmrE homodimer. Notably, interhelical loop residue D84 and C-terminus residue R106, which are hypothesized to have functionally important roles in substrate recognition (Brousseau et al. 2024), both have well-resolved peaks for chains A and B. The same is observed in the 2D ^1^H-^15^N TROSY-HSQC (Fig. 1B) which shows clear resolution between the two chains of the homodimer for nearly every residue of EmrE.

**Figure 1.**
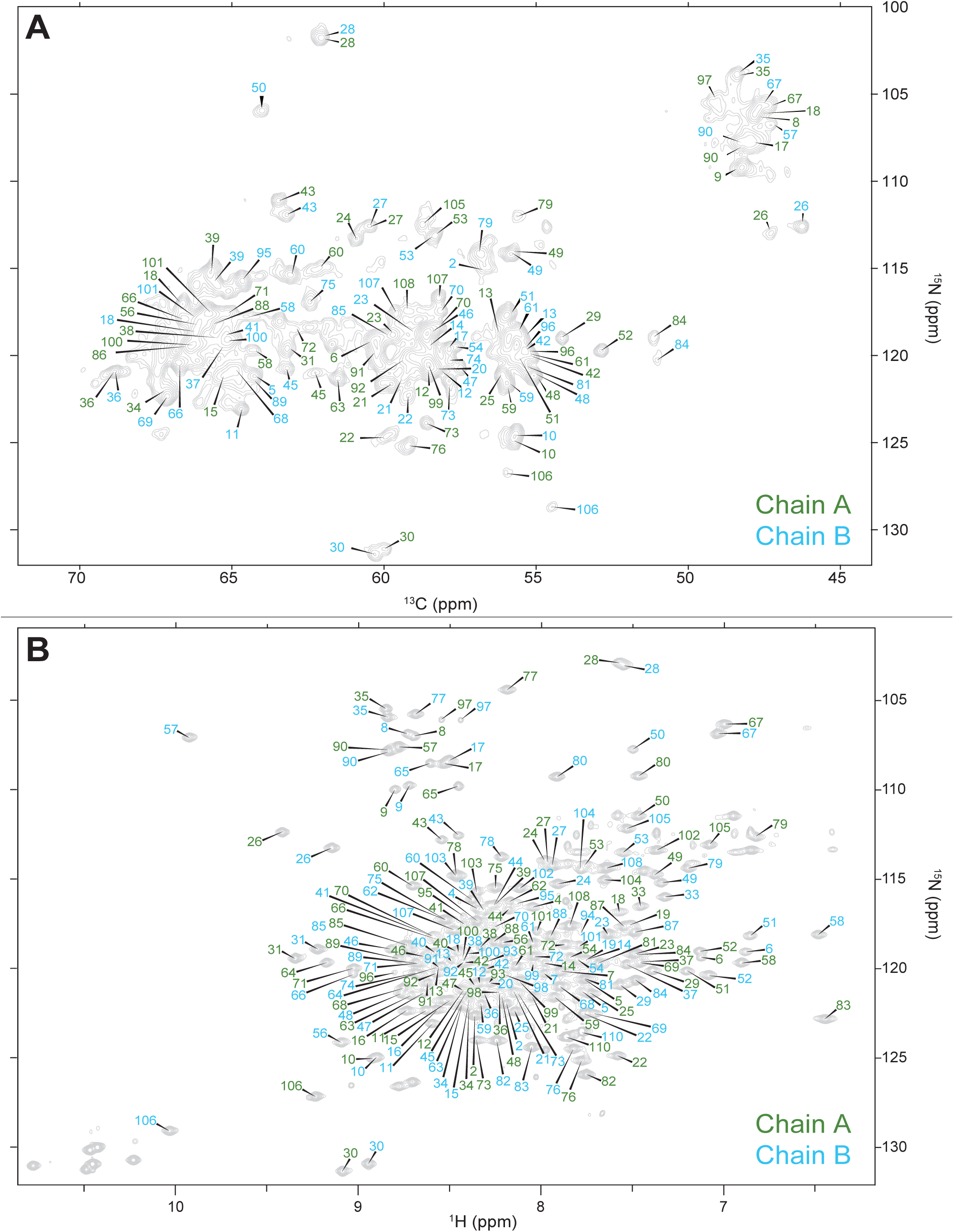
2D fingerprint solid-state (NCA) and solution state (^1^H-^15^N TROSY-HSQC) NMR spectra show well resolved peaks corresponding to each chain of S64V-EmrE.

The backbone walk for a representative stretch of an interhelical loop and a transmembrane helix are shown for both solid-state (Fig. 2) and solution NMR assignments (Fig. 3). The interhelical loop backbone walk establishes backbone connectivity between R29-T28-F27-G26-E25 and the helical backbone walk connects residues I11-A10-G9-G8-L7. Complete (100%) solution NMR backbone ^1^H, ^15^N, ^13^CA, ^13^CB, and ^13^C’ assignments of were carried out with HNCA (red) and HNcaCB (orange) spectra using standard solution backbone walk procedures (Michael Sattler 1999) and are consistent with the chemical shifts of protonated S64V-EmrE in bicelles at pH 5.0 (BMRB ID 51024) that were previously published (Spreacker et al. 2022). The solution assignments are also transferrable and used to guide backbone assignments in the SSNMR spectra. SSNMR backbone assignments were performed at 1.1 GHz using NCOCA, NCACO, and CANCO spectra.

**Figure 2.**
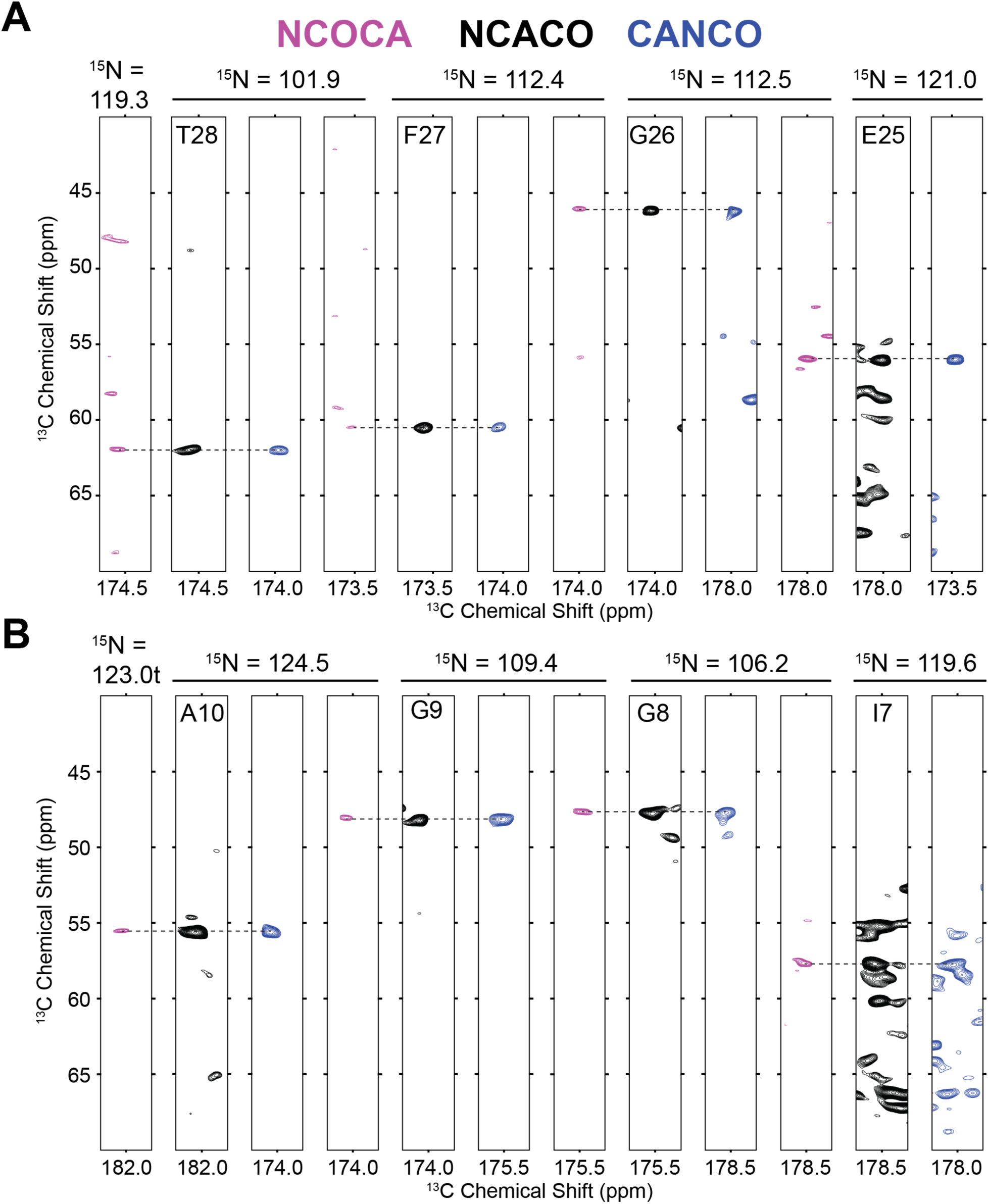
Representative backbone walk for solid state assignment of (A) interhelical loop region E25-R29 and (B) helical region L7-I11 performed using NCOCA (magenta), NCACO (black) and CANCO (blue) NMR spectra.

**Figure 3.**
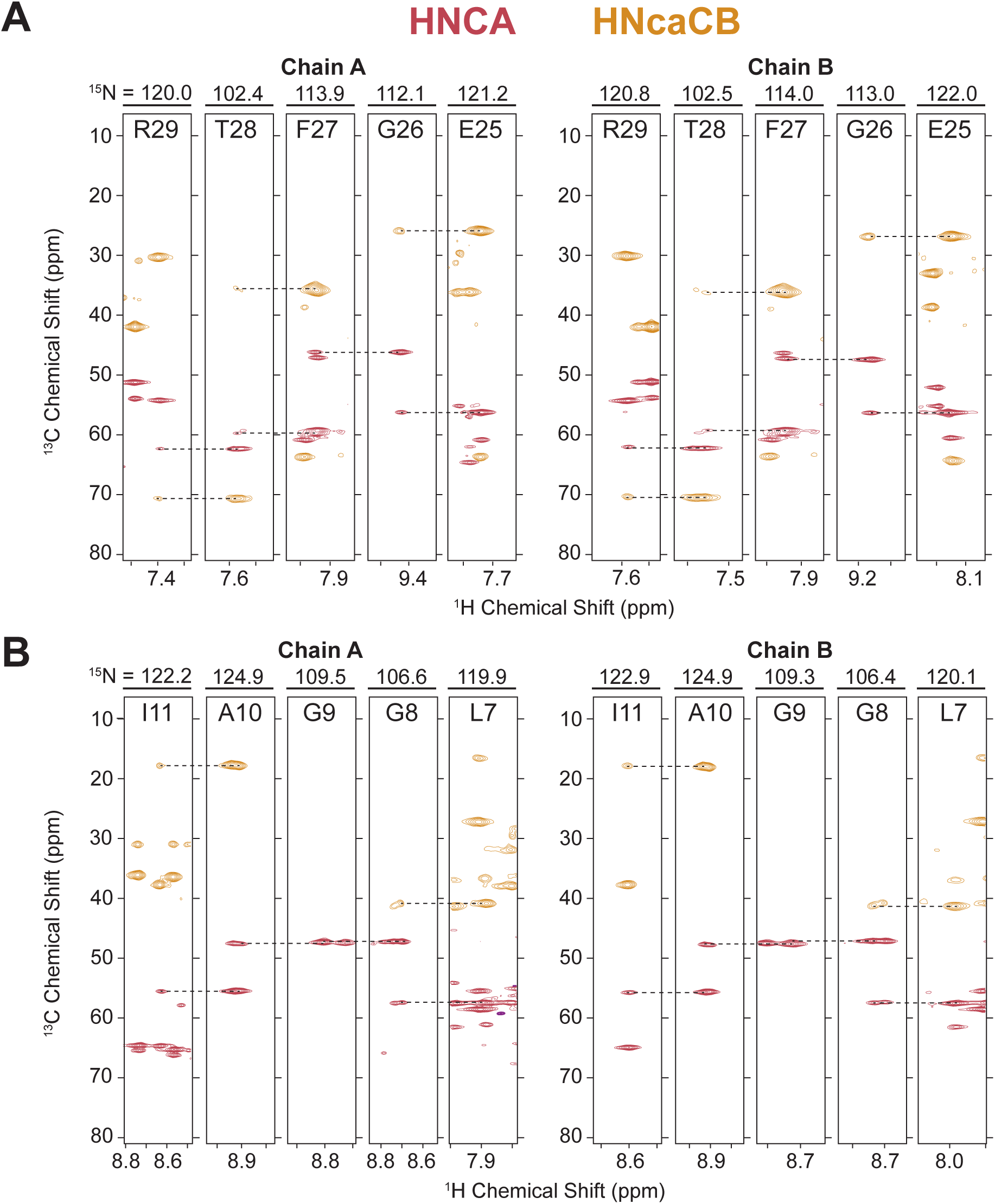
Representative backbone walk for solution NMR assignment of (A) interhelical loop region E25-R29 and (B) helical region L7-Y11 showing inter-residue connectivity through the HNCA (red) and HNcaCB (orange).

### Resonance assignment of isoleucine δ1, leucine, valine, and threonine methyl groups

Complete (100%) of Ile C81, Leu C81 & C82, Val Cγ1 & Cγ2, Ala Cβ, and Thr Cγ2 methyl group assignments of bicelle-reconstituted pH 5.8 S64V-EmrE were carried out through a combination of assignment strategies. These included methyl TOCSY experiment, NOESY experiments correlating methyl resonances to previously assigned backbone amides, and iterative structure-based assignment using methyl NOESY experiments.

#### Matching geminal Leucine and Valine methyl ^13^C resonances using short-mixing time 3D CH_3_-CH_3_ NOESY

We first acquired a 3D methyl NOESY experiment (Catherine Zwahlen 1998) with a 50 ms mixing time on a sample of S64V-EmrE with ^13^C-labelled Cγ1/Cγ2 valine and Cδ1/Cδ2 Leucine. This experiment produced NOESY cross-peaks between the geminal methyl groups of each valine or leucine residue, and these are the dominant NOEs in the spectrum. This allowed geminal pairs to be assigned to the same pseudo-residue.

#### Valine methyl assignments using an HCCH_3_-TOCSY experiment

A 3D HCCH_3_-TOCSY experiment (Uhrin et al. 2000) was acquired to link valine methyl resonances to their corresponding CB and CA chemical shifts. This enabled unambiguous assignment of all valine methyl groups by comparison of the CB and CA chemical shifts in this experiment with the valine CB and CA assignments from the backbone walk experiments. This process also distinguished valine methyl peaks from the leucine methyl peaks in the same region of the ^1^H-^13^C HMQC spectrum.

#### Methyl assignments using Amide-Methyl NOESY experiments

Previously assigned backbone amide resonances for isoleucine, leucine, and valine were linked to their corresponding methyl resonances using two 3D-NOESY experiments in a similar process to (Kurauskas et al. 2017) (Fig. 4). We first used the ^15^N-TROSY-HMQC-NOESY-^13^C-HMQC experiment to match backbone ^15^N and ^1^H resonances to their corresponding methyl carbon resonances. We then navigated to the amide ^15^N and methyl ^13^C plane in the ^13^C-HMQC-NOESY-^15^N-TROSY-HMQC experiment to identify the corresponding methyl ^1^H chemical shifts and assign the methyl peaks in the 2D ^13^C-HMQC spectrum. This approach worked well for assigning peaks with well-resolved methyl resonances, but resonances in areas with high spectral overlap were given ambiguous assignments. Stereospecific Leu and Val assignments were later made using the ^15^N-TROSY-HMQC-NOESY-^13^C-HMQC experiment collected at 1.1 GHz by assigning Leu Cδ2 or Val Cγ2 in accordance with previously published methods. (Mei Hong 2009)

**Figure 4.**
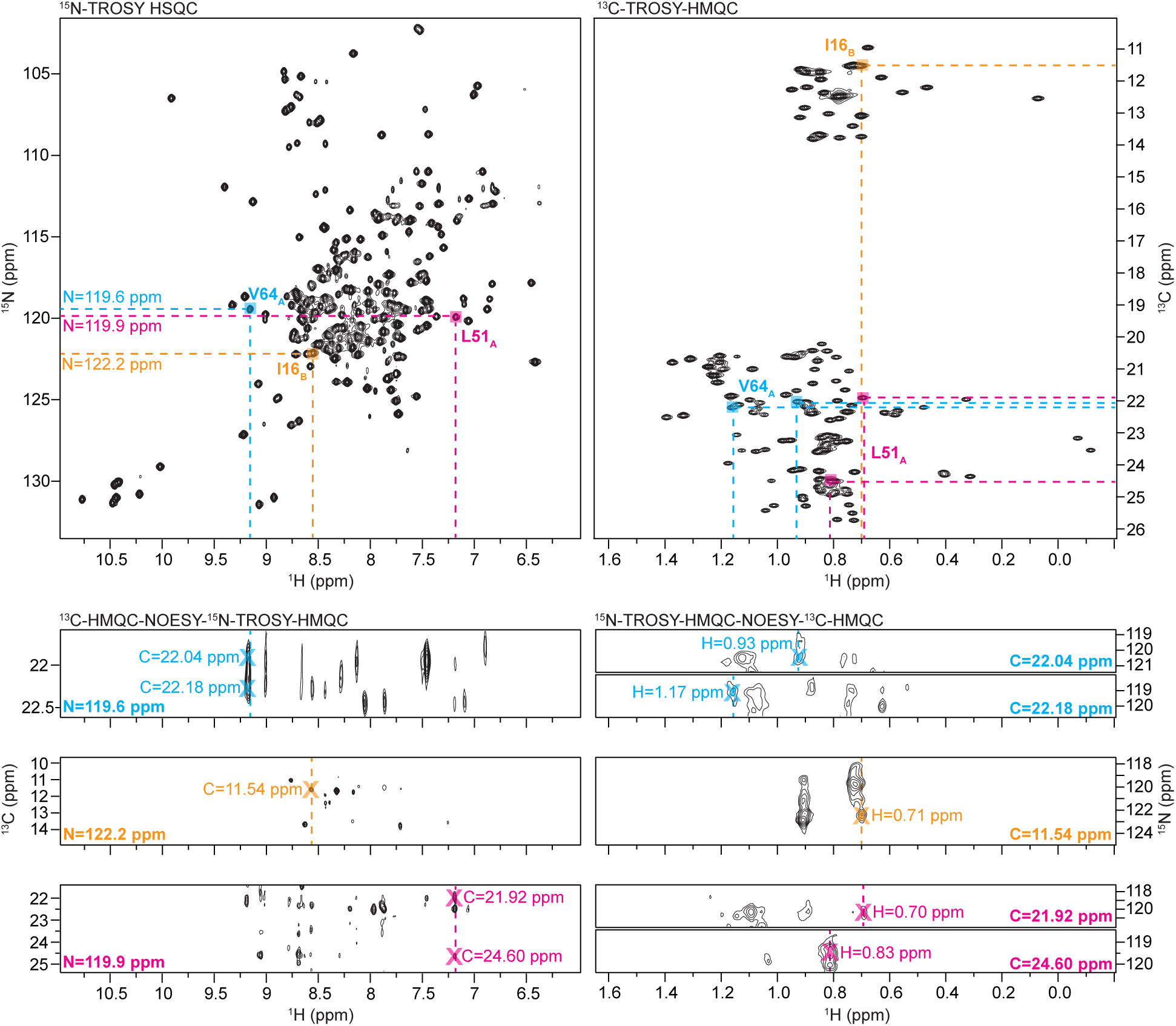
Process of assigning methyl resonances using amide-methyl NOESY experiments: Assigned backbone amide resonances in the 2D ^15^N-TROSY HSQC (top left) are matched to methyl ^13^C resonances in the 3D ^13^C-HMQC-NOESY-15N-TROSY-HMQC (bottom left). Methyl ^13^C resonances are then traced to their corresponding methyl ^1^H resonances in the ^15^N-TROSY-HMQC-NOESY-^13^C-HMQC (bottom right) and matched to peaks in the 2D ^13^C-TROSY-HMQC (top right).

#### Methyl assignments using structural models and ^13^CH_3_-^13^CH_3_ NOESY experiments

The highest-resolution low-pH EmrE crystal structure (Kermani et al. 2022) (PDB 7MH6) and computational structures (Vermaas et al. 2018) (faRM) were used in conjunction with methyl-NOESY experiments and previously assigned methyl resonances to make logical assignments of the remaining methyl groups. Distances between methyl groups were extracted from each structural model using the web-based program NOESY Neighbors (Sung 2022) and in-house scripts. Distances which were within 10 Å and consistent between both structural models were deemed “predicted NOEs.” These predicted NOEs were compared with NOE cross-peaks from the long-mixing-time 3D- and 4D-methyl-NOESY NMR experiments and matches between predicted and experimental NOEs were iteratively assigned through structure refinement and process of elimination.

### Comparison between Solid and Solution NMR Assignments

Backbone (^13^CA, ^13^C’, and ^15^N) and the sidechain ^13^CB chemical shifts between solid and solution state NMR were quantitatively compared for chains A and B (Figure 5). The ^13^CA, ^13^C’, ^15^N’ and ^13^CB chemical shifts have Pearson correlation scores of 1.00, 0.99, 0.99, and 1.00, respectively, for both chains in the asymmetric homodimer. The agreement between solid and solution-state chemical shifts is indicative of consistent secondary structure between EmrE in q=0.33 DMPC/DHPC isotropic bicelles and in POPC bilayers. We also observe close agreement for backbone and CB chemical shifts between solution NMR data acquired above the phase transition and solid-state NMR data acquired below the phase transition temperature, enabling assignments of sidechain residues in the low temperature ^13^C-^13^C, ^15^N-^13^C-^13^C, and ^13^C-^13^C-^13^C solid-state NMR experiments.

**Figure 5.**
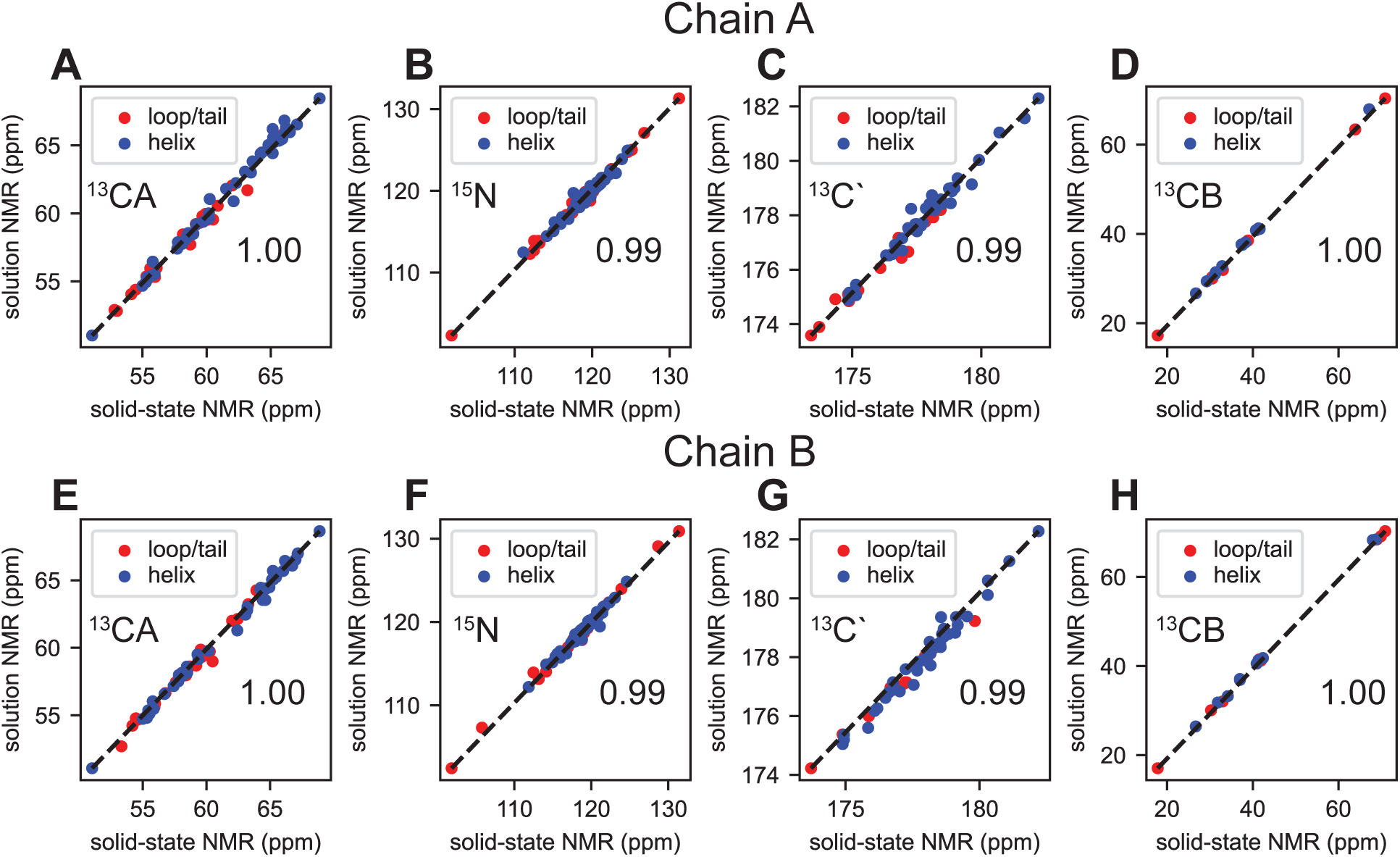
Correlation plots comparing backbone ^13^CA, ^15^N, ^13^C’, and sidechain ^13^CB chemical shifts from solid state and solution state NMR backbone experiments for chain A (A-D) and chain B (E-H). Residues which exhibit helical properties are displayed in blue and residues which are in the loops or C-terminus of EmrE are shown in red. Solid state chemical shifts are using spectra collected at 1.1 GHz and above the phase transition. The number of points in each plot for chain A are (A) 70, (B) 65, (C) 59, and (D) 19 for chain A. The number of points in each plot for chain B are (E) 65, (F) 64, (G) 54, and (H) 21. The Pearson correlation coefficients are (A) 1.00, (B) 0.99, (C) 0.99, (D) 1.00, (E) 1.00, (F) 0.99, (G) 0.99, and (H) 1.00 and illustrate agreement between solution and solid-state NMR chemical shifts.

### Sidechain Assignments using Solid-State NMR

A total of 65% of the non-aromatic ^13^C sidechain atoms in EmrE have been assigned using NCACX, NCOCX, ^13^C-^13^C, and ^13^C-^13^C-^13^C SSNMR spectra and building off the backbone assignments. The ^13^C-^13^C spectrum is a foundational fingerprint spectrum and serves as a roadmap for identifying outlying spin systems in chemically unique environments (Fig 6). Some examples include E25, T28, A52, and D84, all of which are in located in interhelical loops. While chemical shifts of E25, T28, and A52 are observed to be closely overlapped for both chains, D84 is not. Moreover, D84_A_ CA and CB crosspeaks are more intense than those of D84_B_; and D84_A_ CG and CD crosspeaks are observed, while the those in D84_B_ are not.

**Figure 6.**
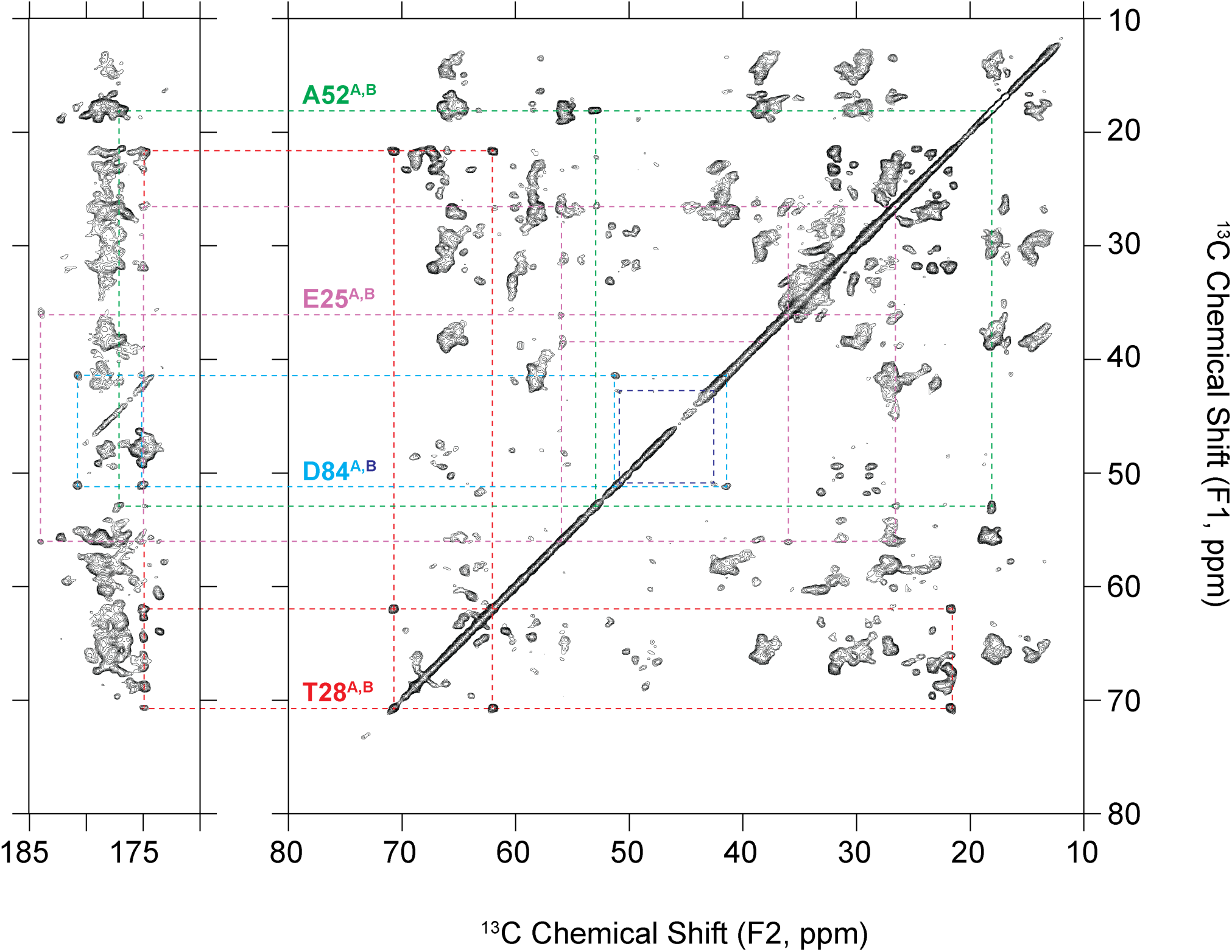
Representative spin systems assigned in EmrE in a ^13^C-^13^C (DARR, t_mix_ = 50 ms) correlation. Each spin system is isolated and in a unique chemical environment because they exist in interhelical loops.

### Conclusion

Solution- and solid-state NMR assignments are reported for S64V-EmrE at pH 5.8. These include ^1^H, ^15^N, and ^13^C resonance assignments of the backbone from both solution- and solid-state NMR, Ile C81, Leu C81 & C82, Val Cγ1 & Cγ2, Ala Cβ, and Thr Cγ2 methyl groups from solution NMR experiments, and side-chain assignments from solid-state NMR ^13^C-detected experiments. The backbone assignments are consistent between solution- and solid-state NMR. Solution- and solid-state NMR resonance assignments will be used to calculate a high-resolution structure of S64V-EmrE.

## Author Contributions

AH prepared both solid and solution NMR samples of EmrE. BDH, SW, and CMR collected and processed SSNMR spectra and AH, MT, and KHW collected and processed solution NMR spectra. BDH and CMR assigned SSNMR spectra and AH and KHW assigned solution EmrE spectra. BDH, AH, CMR, and KHW wrote the manuscript.

## Data Availability

Solid-state NMR resonance assignments of S64V-EmrE in liposomes are available through the BMRB accession number 52793. Datasets collected at 1.1 GHz are publicly available on the Network for Advanced NMR (NAN) Data Browser under ARK IDs ‘66276e0a-fdca-443e-bdd5-bfcdcdbf22a9.1’ (CANCO), ‘9bbc0d40-3f21-453e-bb11-39452d726f03.1’ (NCACO) and ‘f8b8e168-50ea-4f5a-93d7-9f101630b97a.1’ (NCOCA) and all other data is available under the BMRBig ID ‘bmrbig110’. Reported solution-state NMR resonance assignments of S64V-EmrE in isotropic bicelles are available through the BMRB accession number 52790 and datasets for solution-state NMR experiments are available on the Network for Advanced NAN Data Browser under ARK ID ‘c5690d6c-b839-4b37-82a4-393e903501e3.2’ and BMRbig under ID ‘bmrbig109’.

## Declarations

Authors declare no competing interests

